# Whole Genome Detection of Sequence and Structural Polymorphism in Six Diverse Horses

**DOI:** 10.1101/545111

**Authors:** Mohammed Ali Al Abri, Heather Marie Holl, Sara E Kalla, Nate Sutter, Samantha Brooks

## Abstract

The domesticated horse has played a unique role in human history, serving not just as a source of animal protein, but also as a catalyst for long-distance migration and military conquest. As a result, the horse developed unique physiological adaptations to meet the demands of both their climatic environment and their relationship with man. Completed in 2009, the first domesticated horse reference genome assembly (EquCab 2.0) produced most of the publicly available genetic variations annotations in this species. Yet, there are around 400 geographically and physiologically diverse breeds of horse. To enrich the current collection of genetic variants in the horse, we sequenced whole genomes from six horses of six different breeds: an American Miniature, a Percheron, an Arabian, a Mangalarga Marchador, a Native Mongolian Chakouyi, and a Tennessee Walking Horse. Aside from extreme contrasts in body size, these breeds originate from diverse global locations and each possess unique adaptive physiology. A total of 1.3 billion reads were generated for the six horses with coverage between 15x to 24x per horse. After applying rigorous filtration, we identified and functionally annotated 8,128,658 Single Nucleotide Polymorphisms (SNPs), and 830,370 Insertions/Deletions (INDELs), as well as novel Copy Number Variations (CNVs) and Structural Variations (SVs). Our results revealed putatively functional variants including genes associated with size variation like *ANKRD1* and *HMGA1* in the very large Percheron and the *ZFAT* gene in the American Miniature horse. We detected a copy number gain in the *Latherin* gene that may be the result of evolutionary selection for thermoregulation by sweating, an important component of athleticism and heat tolerance. The newly discovered variants were formatted into user-friendly browser tracks and will provide a foundational database for future studies of the genetic underpinnings of diverse phenotypes within the horse.

**Author Summary:** The domesticated horse played a unique role in human history, serving not just as a source of dietary animal protein, but also as a catalyst for long-distance migration and military conquest. As a result, the horse developed unique physiological adaptations to meet the demands of both their climatic environment and their relationship with man. Although the completion of the horse reference genome yielded the discovery of many genetic variants, the remarkable diversity across breeds of horse calls for additional effort in quantification of the breadth of genetic polymorphism within this unique species. Here, we present genome re-sequencing and variant detection analysis for six horses belonging to geographically and physiologically diverse breeds. We identified and annotated not just single nucleotide polymorphisms (SNPs), but also large insertions and deletions (INDELs), copy number variations (CNVs) and structural variations (SVs). Our results illustrate novel sources of polymorphism and highlight potentially impactful variations for phenotypes of body size and conformation. We also detected a copy number gain in the *Latherin* gene that could be the result of an evolutionary selection for thermoregulation through sweating. Our newly discovered variants were formatted into easy-to-use tracks that can be easily accessed by researchers around the globe.

## Introduction

Quantifying genetic variation is an important theme in modern biology and population genetics. Recent technological advances in genomics have benefitted livestock by allowing examination of genetic variation in these non-traditional model species at an unprecedented scale and resolution. Cataloging that variation lays the foundation for dissecting the complex genetic architecture of phenotypic variation, which in turn has many applications in livestock health, welfare, physiology and production traits [1,2]. Application of these variations also improves inference of ancient demographic and evolutionary histories, as well as the mechanisms underlying adaptation in various species [3]. In addition, cross-species comparisons of genetic variation improves our understanding of the structure-to-function relationship within conserved elements of the mammalian genome [4].

Domesticated approximately 5,500 years ago, horses were historically used for food, transportation, trade, warfare, and as draught animals [5]. Man has since selected horses suitable for a range of physical and behaviorally desirable traits, ultimately resulting in the formation of more than 400 unique horse breeds [6]. Comparisons between ancient and domesticated horse genomes revealed signatures of this selective pressure in ~125 potential domestication target genes [5]. Advantageously, the *Equidae* possess a particularly old and diverse fossil record, aiding not only in characterizing their demographic history but also ancient human movement and migration [3,5,6]. However, compared to other livestock species, relatively few studies have focused on the discovery of the standing genetic variation within different horse breeds [7]. Therefore, additional investigation of the equine genomic architecture is critical for a better understanding of the equine genome, as well as for expanded comparisons across diverse mammalian species. Furthermore, the equine industry itself provides an eager opportunity to apply genomic discoveries towards improvements in the health and well-being of this valuable livestock species.

Here, we sequenced six horses belonging to six divergent breeds in order to enrich the current collection of genetic variants in the horse. Namely, we sequenced a female Percheron (PER) and an American Miniature (AMH), and a Tennessee Walking Horse (TWH), a Mangalarga Marchador (MM) and a Native Mongolian Chakouyi (CH) male horses. These breeds were historically selected to perform distinct tasks and therefore may harbor a wealth of unique variation at the genome level. For example, the Arabian horse was primarily selected for metabolic efficiency, endurance and strength. Severe desert conditions required a resilient animal with absolute loyalty. Arabian horses were central to the survival and culture of the Bedouins peoples of the Arabian Peninsula [8]. The Percheron horse was primarily selected for large size and developed as draft horse and was used as both a war horse and a farm horse [9]. On the other hand, the American miniature horse was selected for small body size [10,11]. The Mangalarga Marchador is a Brazilian Iberian horse breed that was selected for its versatility and stamina. It is capable of performing diverse tasks, from performance racing to herding cattle [12]. The Tennessee Walking Horse (or Tennessee Walker) is a unique in that it is the only U.S. breed able to perform an even-timed 4-beat gait called the “running-walk” at intermediate speeds [13].

Our goal was to investigate not just single nucleotide polymorphisms (SNPs) and small insertion deletions (INDELs), but also copy number variations (CNVs) and structural variations (SV). After applying rigorous filtration criteria to the read qualities, we detected and annotated the variation in each of these four classes within the six horses. These genetic variants are now publically available in European Nucleotide Archive (ENA) as well as in the National Animal Genome Research Program (NAGRP) Data Repository. CNVs and SVs are often difficult to access in public databases, therefore we have processed these novel variants into user-friendly tracks available for download at https://www.animalgenome.org/repository/horse/6_horse-breeds_variants/.

## Results and Discussion

### Whole genome sequencing, alignment and quality control

High throughput Next Generation Sequencing (NGS) provides affordable access to the enormous collection of existing genetic variation in genomes. We used paired-end Illumina sequencing to interrogate the genomes of an Arabian, a Percheron (a breed of primarily French origin), an American Miniature, a Mangalarga Marchador (from Brazil), a Native Mongolian Chakouyi, and a Tennessee Walking Horse to an average sequence coverage of 10x to 24x. Sequencing was completed using the Illumina HiSeq2500 (Illumina, San Diego, CA) with manufacturer recommended reagents and procedures by the Biotechnology Resource Center at Cornell University. A total of 1.3 billion reads were generated for the six horses. The raw number of reads ranged between 142 million reads on the Native Mongolian horse to 324 million reads on the American miniature (**Table 1**). After filtering, between 83 million (Mangalarga Marchadore) and 187 million (Percheron) paired reads were retained. Both read pairs aligned to the EquCab2 reference genome [1] in 92 - 97 % of reads, indicating a successful mapping procedure (**Table 1**).

**Table 1:** Yield, filtering and mapping summary of the next generation sequencing data of six horses from different breeds. The depth of coverage and mapping metrics show a descent quality of the 6 genomes sequencing and genome coverage.

|  | ARB | PER | AMH | TWH | MM | CH |
| --- | --- | --- | --- | --- | --- | --- |
| Number of paired-end reads before trimming | 241,480,555 | 296,460,133 | 324,123,384 | 198,749,393 | 169,680,137 | 142,502,233 |
| Read lengths | 100/100 | 100/100 | 100/100 | 150/150 | 150/150 | 150/150 |
| Estimated average depth of coverage before trimming<sup>1</sup> | 17.8x | 21.96x | 24x | 22.0x | 18.9x | 15.833x |
| Number of reads after trimming | 165,277,009 | 187,223,705 | 138,772,441 | 161,659,278 | 134,732,394 | 121,744,242 |
| Total number of aligned reads | 330,554,018 | 374,447,410 | 277,544,882 | 323,318,556 | 269,464,788 | 243,488,484 |
| Estimated depth of coverage<sup>1</sup> | 12.24x | 13.87x | 10.28x | 17.96x | 14.97x | 13.53x |
| Percentage of mapped reads<sup>2</sup> | 98% | 93% | 93% | 97% | 95% | 96% |
| Percentage of reads where both pairs mapped<sup>2</sup> | 97% | 92% | 92% | 96% | 94% | 94% |

1. Estimated from trimmed and aligned reads using the formula $C = L * N / G$ (where G is the haploid genome length ( $2.7 * 10^9$ ), L is the read length and N is the number of reads) 2. Estimated using the bamtools *stats* procedure in bamtools.

### Identification of Variants

#### SNPs

In total, 8,562,696 SNPs were detected using the GATK *HaplotypeCaller* and processed using the GATK *VariantRecalibrator* procedure, producing a final set of 8,128,658 SNPs. The number of SNPs is about 0.3 % of the size of the genome (about 1 every 300 base pairs) which is very similar to the percentage of SNPs estimated in the human genome [14]. Amongst those, 11,537 SNPs (0.14 %) were multi-allelic. The mean transition to transversion ratio in these horses is 1.998 (range 1.991 to 2.008) (**Table 2**), a value very close to other mammalian species [15].

**Table 2:**
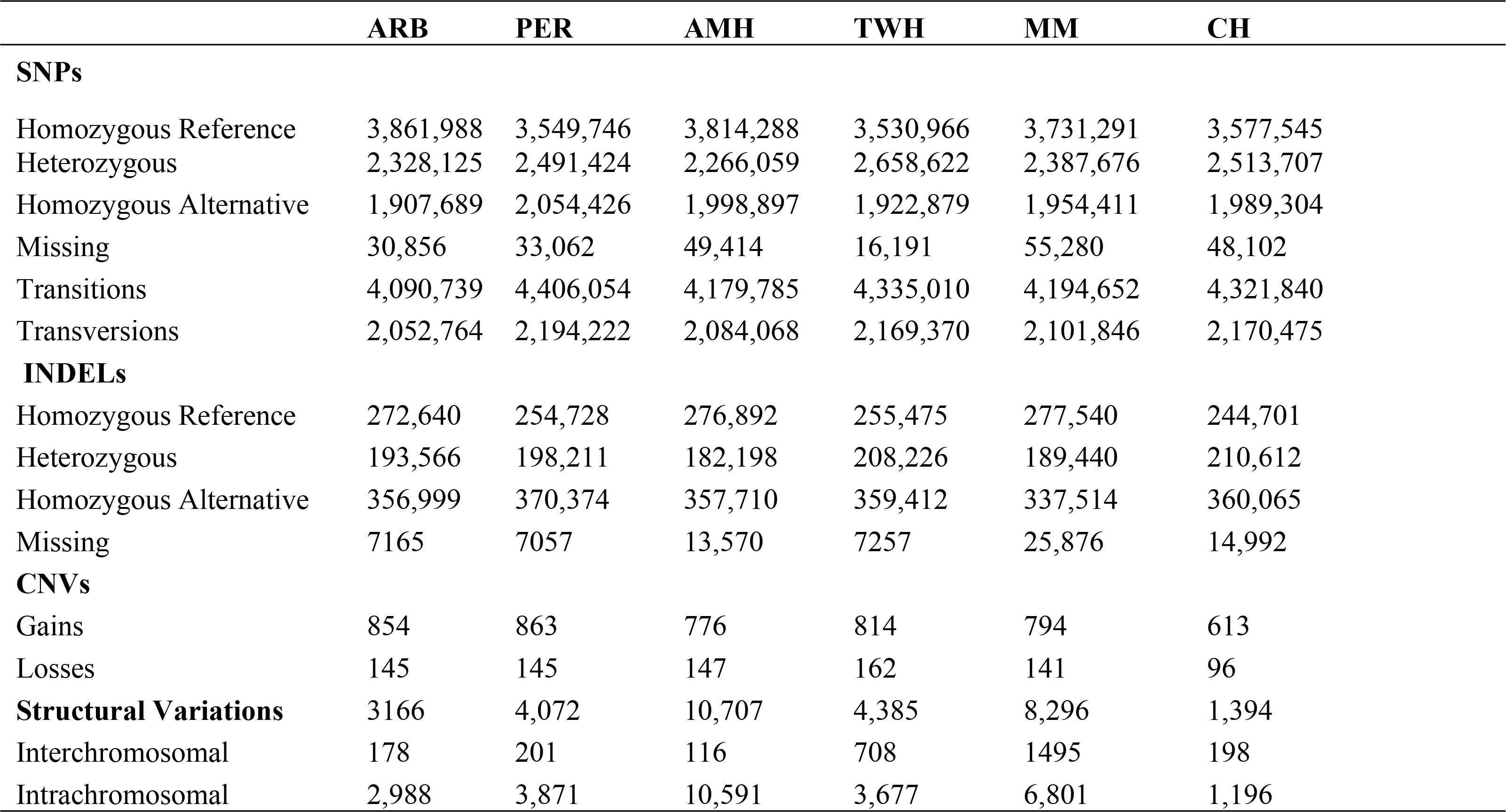
Genotype categories of SNPs and INDELs and counts of CNVs and SVs in the six horses.

The allelic frequency spectrum (**Figure S1**) showed an expected decline in the frequency of SNPs as the observed number of the alternative allele increased, as observed in other studies [16],[17]. The mean, median and standard deviation of Phred-scaled quality scores for the SNPs were 785.78, 543 and 732.85, respectively, which signifies a very high call accuracy.

Two of the sequenced horses were genotyped previously using the Illumina EquineSNP50 array (Illumina Inc.) enabling a test of the genotype concordance between the array and sequencing methods. The concordance of the genotypes detected using the Equine SNP50 genotyping array and those detected by NGS was 96% for the American Miniature and 98% for the Percheron horse, illustrating that the SNPs detection is comparable to array-based methods and is reliable for the purposes of this study.

The highest proportion of genotype calls were the homozygous reference genotype, comprising 43 to 47 % of the SNPs in each horse (**Table 2**). Relative to the chromosome size, the highest proportion of SNPs was found in chromosome 12 (0.5 %) followed by chromosome 20 (0.4%) (**Figure 1**). Overall, the highest proportion of homozygous reference genotypes was found in the Arabian horse. This may be explained by the fact that, among the breeds included, the Arabian horse has the closest historical relationship to the reference genome derived from a mare of the Thoroughbred breed. The Thoroughbred horse population originated by mating three prominent Arabian stallions to native mares in England during the 17^th^ century [18].

**Figure 1.**
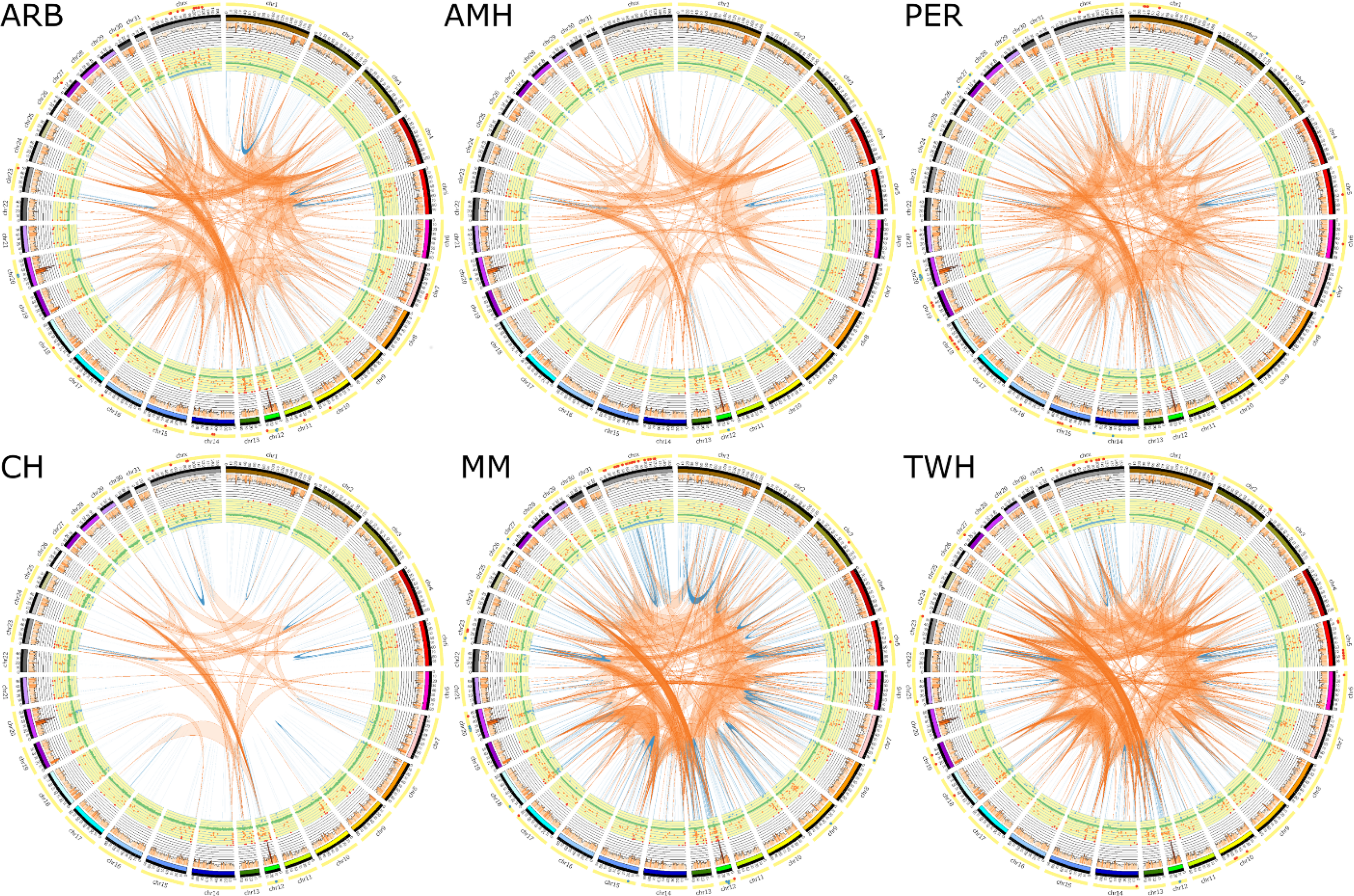
Circos plot summarizing the genetic variants detected in each horse. The pattern of variation across the genomes of the six horses reveals structurally diverse regions important for immunity and olfactory reception at chromosomes 12 and 20 in all six horses. From the inside out, each plot shows two endpoints of the inter-(orange) and intra-(blue) chromosomal translocations. Intrachromosomal translocations > 5 MB are in dark blue. The yellow ring shows the copy number variations (green =normal, blue = loss, red=gain). The histogram (in orange) shows the density of SNPs detected using 1MB windows. The outermost track in yellow marks the lower 1% (red) and upper 1% (blue) values of the average nucleotide diversity (π) calculated using 1 MB windows.

#### INDELs

It is well established that INDELs are the second most common form of genomic variations, altering a similar total proportion of base pairs as SNPs [19]. In horses, some known INDELs cause genetic disorders such as the Lavender Foal Syndrome (LFS) [20] and the Severe Combined Immunodeficiency (SCID) [21]. We detected 830,370 small INDEL loci using the GATK *HaplotypeCaller* procedure. Within this set, 10,811 INDELs were multi-allelic. The mean, median and standard deviation of Phred-scaled quality scores for the INDELs were 1,025, 785 and 1,076 respectively, which signifies a higher accuracy but greater dispersion in accuracy, to that observed in SNPs. INDEL size ranged from 2 to 233bp, with mean of 39 bp and a median of 83 bp. The INDELs were split almost equally between insertions and deletions (48 % and 52 % respectively), as was observed in the INDELs pattern in humans [22], [23]. Unlike SNPs, the most frequent small INDELs calls were the homozygous alternative calls, ranging between 40 and 44 % of the total INDELs calls in each horse (**Table 2**). INDELs are more rare events than SNPs and are thus more likely to be unique to a breed of horses than to be shared between breeds [24]. In fact, the resolution of Eukaryotic phylogenetic trees can be improved by incorporating INDELs [25].

It is noteworthy that the incidence of homozygous alternative (non-reference homozygous) SNPs and INDELs was highest in the Percheron horse. This could be a result of a greater divergence between the reference genome (a Thoroughbred horse) and the Percheron relative to the other breeds. The SNP and INDEL genotype missingness was the highest in the Mangalarga Marchadore and was inversely correlated with the aligned reads depth (before trimming) (**Tables 1 and 2**). The total number of INDELs called within each genome was influenced by the depth of coverage, as previously observed in NGS data of human genomes [26].

#### CNVs and SVs

CNVs and SVs contribute significantly to genomic diversity [27]. Genome-wide datasets produced by NGS technologies are revealing a wealth of knowledge about the frequency and structure of these types of polymorphisms. Of the identified CNVs, the number of gains was consistently higher than the number of losses for all horses (**Table 2**, 786 mean gains vs 139 mean losses). Since many of the gains are shared between horses, we hypothesize that the excess of gains is an artifact of the computational assembly of EquCab2.0, compressing regions of repetitive sequences and highly homologous gene families. Numerous CNV regions (or genes) in the genome could represent duplication events incorrectly assigned to regions of high homology in EquCab2. Some of those regions were annotated within the unscaffolded “ChrUn”, which rendered them difficult to study, and likely to represent expanded gene families and repetitive elements unamenable to computational assembly.

Additionally, we also observe a consistent excess of intrachromosomal SVs compared to the interchromosomal SVs (**Table 2**). Bias towards intrachromosomal SVs is not uncommon in this type of analysis and is often due to a preference for intrachromosomal joining resulting from the relative closer proximity of these genomic regions. This same phenomenon was observed in studies of the mouse [28], human [29] and chicken [30]. It is proposed that a biological mechanism preferring proximal intrachromosomal rearrangement reduces large-scale genomic alterations, and therefore maintains genomic stability [28].

The American miniature horse possessed the highest number of the Intrachromosomal SVs (n= 10,591) compared to other horses. These results are more than double the average of the corresponding values in the other horses and may be an artifact of imperfect library preparation or fragment size selection prior to sequencing. Indeed, filtering of such artifacts is a significant challenge for reliable discovery and genotyping of SVs by sequence based methods. Notably, more interchromosomal SVs were detected in the Mangalarga Marchador and the Tennessee Walking Horse (n=1,495 and n=708 respectively). Sequencing libraries from these horses also possessed the largest average insert size (248 bp and 207 bp respectively). Larger insert sizes allow for the discovery of relatively more SVs in NGS data [31] and comparisons of SVs totals across individuals with varying insert size need to account for this effect.

### Annotation of Detected Variants

The majority of SNPs were intergenic, followed by intronic, comprising 60 % and 27 % of SNPs, respectively (**Table S1**) [32]. The small percentage of exonic SNPs (0.08 %) and INDELs (0.06%) is likely the result of strong negative selective pressure exerted on coding regions due to the potentially severe functional implications of these alterations [23]. Likewise, a lower diversity around 3’ UTR and 5’ UTR regions was found in SNPs and INDELs (0.02 % - 0.07 %), which was also reported in other mammalian species [23,32] (**Table S1**). In terms of predicted functional impact, the majority of SNPs found in coding sequences (33,593, 0.39%) were synonymous (**Table S1**). Yet, 26,322 (0.30 %) SNPs were predicted to result in a non-synonymous amino acid change and 2,863 (0.316%) INDELs caused frame-shifts. The highest concentration of SNPs in the genomes of all the horses was observed amidst ECA12 and ECA20. Functional annotation of these regions showed that they are involved in metabolic and sensory perception (ECA12) and immune response and antigen processing (ECA20). This could be indicative of an evolution of these genes. However, it might be the result of mis-assembles in the reference genome or misalignment of reads (which is unlikely to occur in all six genomes simultaneously).

Copy number and structural variations are given relatively little attention compared to SNPs in studies of genetic diversity. Nevertheless, they are ubiquitous in the mammalian genome and influence a number of phenotypes [33,34]. The resulting CNVs and SVs were annotated using the ENSEMBL genes they overlap with and are available at (https://www.animalgenome.org/repository/horse/6_horse-breeds_variants/). We found that chromosomes ECA12 and ECA20 possessed the highest density of CNVs. Functional annotation of these regions revealed large gene families involved in olfactory reception and immunity. The same observation was made by array CGH in the horse [33] and corresponding findings were also obtained in the human genome [35].

Our functional CNVs annotation revealed a copy number gain in a gene cluster that includes Latherin (*LATH*) (ENSEMBL Gene Symbol: ENSECAG00000009747.1). The reference genome assembly indicates just one copy of this gene, yet a copy-number gain was previously reported in a Quarter Horse using NGS data [7], and using array CGH a copy number loss was observed in the same region [33] (ECA 22:23893194-24009882). *LATH* (also known as *BPIFA4*) is a member of the palate lung and nasal epithelium clone (*PLUNC*) family of proteins that is common in the oral cavity and saliva of mammals [36,37]. Based on ENSEMBL gene models (V.86) and alignment of non-horse RefSeq genes (UCSC), the region surrounding equine *LATH* in EquCab2.0 contained evidence for 14 BPIF family gene models. In contrast, the orthologous bovine region in BosTau8 contained just ten, and the human genome (hg38) has eight BPIF RefSeq gene models. In horses this gene produces a surfactant protein that is expressed in the saliva and uniquely to the *Equidae*, as the primary protein component of sweat [38]. Therefore, equine Latherin protein may play an important role in mastication of fibrous food, as well as in enhancing evaporative cooling [36]. Therefore, it is reasonable to postulate that the gain in *LATH* copies observed in this study results from an evolutionary pressure for improved evaporative dissipation of heat, yielding athleticism and endurance in hot environments.

The CNV gains/losses across the *LATH* regions were common across the six horses included in the study, suggesting that the reference genome may possess errors in this region. We therefore validated the presence of a copy number variation in the *LATH* gene and genes surrounding it using quantitative PCR (qPCR) analysis in eight horses, including the horse used to produce the EquCab2.0 assembly. QPCR showed evidence of between two and six copies of *LATH* relative to a single copy control gene (*ASIP*) across X individual horses of diverse breeds, although the limited resolution of the qPCR approach could not significantly differentiate copy numbers for individual animals when compared to the horse utilized in the reference genome assembly (**Figure 2)**. Notably, the reference genome animal (“Twilight”) possess a mean copy number of four, by qPCR, though the assembly derived from her documents just one *LATH* locus. On the other hand, the QPCR analysis indicated statistically significant difference in copy number between horses of some nearby genes (*BPIFB4*, and *BPIFA1*) but not *BPIFA2*. Precise haplotype analysis of this complex CNV polymorphism is challenging due to the poor quality of the reference assembly within this region, and the technical challenges of qPCR result in limited resolution for this application. Thus, more precise determination of polymorphic CNVs and gene family expansion in the *LATH* region will require the use of more advanced techniques like long-read sequencing and digital qPCR quantification of the CNV in future work [39].

**Figure 2:**
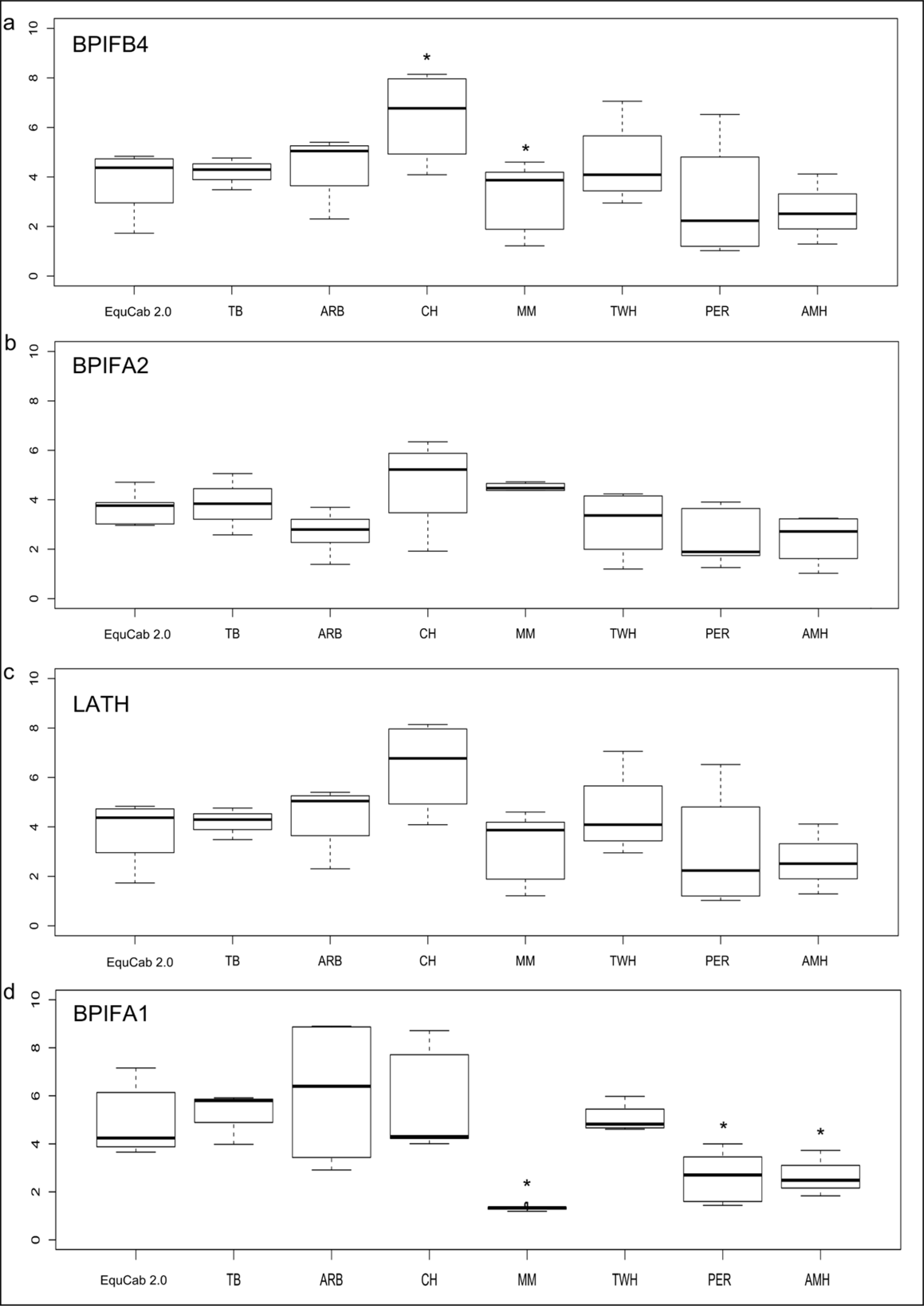
RT-qPCR results of the *LATH* CNV region. Results for different primers are shown relative to their position in the genome. No evidence of a copy number variation is seen for *BPIFB4* (**a**) and *BPIFA1* (**d**) two genes that flank *LATH* (**c**) in the EquCab 2.0 assembly.

Our annotation of the SVs also indicated duplication events within the *ZFAT* gene (ECA 9: 75076827 bp, **Table S2**) unique to the American Miniature horse. The *ZFAT* region is associated with withers height and overall skeletal size in horses [40,41]. We also detected an inverted duplication unique to the Percheron horse in *HMGA1*. In the mouse, a *HMGA1/HMGA2* double knockout produces the “superpygmy” phenotype [42]. Mapping efforts previously identified an effect on overall body size in horses due to the *HMGA2* region [41]. Later, it was shown that a non-synonymous mutation in the first exon of *HMGA2* reduced withers height by an average of 9.5 cm per copy of the mutant allele in small Shetland ponies [43]. These small SVs may impact regulatory motifs in these genes, leading to the size phenotypes observed in these breeds. However, future work is needed to investigate the impact of this new inverted duplication on height or growth within the genetic background of these breeds.

### Genome-wide diversity (π)

Nucleotide diversity (π) [44] is defined as the average number of nucleotide differences per site between two randomly chosen sequences in a population. Assessment of nucleotide diversity provides a valuable insight into the divergence of populations, inferring the demographic history of the species, as well as the historical size of the population [45]. Areas of lower than expected nucleotide diversity may signify signatures of past selection events [46]. Traditionally, such regions are found by comparing the same sequences from multiple individuals [44]. However, π as implemented in VCFtools is calculated from a single genome of a diploid individual [47–49]. In this study, regions with very low diversity were found by calculating π for each horse genome as the average number of differences between two chromosomes using one megabase (MB) non-overlapping windows in VCFtools [50].

The average nucleotide diversity across all six horses was 0.00097 for all SNP polymorphisms, and ranged from a minimum of 0.00090 for the American Miniature Horse and 0.0011 for the Tennessee Walking Horse. This could reflect a higher inbreeding in the American Miniature horse compared to the Tennessee Walking Horse. Average diversity in the autosomal chromosomes for all horses was 0.0010, which is four times as high as the mean diversity observed in the X chromosome (0.00026). Since the X chromosome has three-quarters the effective population size (*N*_e_) of that of the autosomes, lower nucleotide diversity for the X chromosome is to be expected. However, a lower diversity level could also be due to a lower mutation rate (μ) on ECAX. Clearly, X copy number differences specifically influence the calculation of nucleotide diversity levels in the male horses used in this study, as compared to the female reference animal (**Figure 1**).

Notably, the SNP dense regions on ECA20 and ECA12 were amongst the highest (top 1%) in nucleotide diversity (π) value (**Figure 1**). We used PANTHER (v14.0) [54] statistical over-representation test (using a Bonferroni correction at P < 0.05) analysis of xenoRefGene genes in these regions after removing exact duplicate gene names as horses have relatively few refseq genes (**Table S3**). PANTHER further removed duplicate genes, keeping only a single gene ID cases at loci with two or more xenoRefGene names. The analysis revealed that enrichment for the T cell receptor signaling pathway and adaptive immune response on ECA20 and sensory perception on ECA12 (**Table S4**). Paralogous regions like those in large gene families catalyze a collapse of these regions within computational genome assemblies leading to an inflation in the number of SNPs at these loci.

The bottom 1% of the empirical distribution of π values for each horse (**Table S5**) gives potential selected regions. PANTHER statistical over-representation test (using a Bonferroni correction at P < 0.05) of genes in these regions (**Table S6**) revealed regions significantly for response to cold and temperature stimulus in the Arabian horse and genes involved in negative regulation of apoptotic processes and cell differentiation in the Mangalarga Marchador horse. In the Tennessee Walking Horse, the test revealed enrichment for the regulation of cellular processes and inorganic ion homeostasis. In the American Miniature horse, the same test revealed genes involved in development processes and cellular development and in the Percheron horse, it highlighted regions significantly enriched for genes involved in regulation of cell morphogenesis, cytoskeleton organization and actin cytoskeleton organization. These regions included the gene *ANKRD1* which contributes to body size variation in horses [10]. In the Native Mongolian Chakouyi horse, no statistically significant over-represented terms were found. We also investigated the functional classification of gene models from these regions using GO-slim biological processes in PANTHER. Notably, this analysis showed that the Percheron and American Miniature horses were the only two horses with regions containing genes in the category “Growth”, although it formed a small percentage of the total number of genes (**Table S7**).

The low diversity regions found in this study did not overlap with the regions previously reported in Petersen *et al.* using Illumina SNP50 Beadchip and an F_ST_-based statistic [51], likely due to the differences in sample size as well as the methodology. On the other hand, three gene regions reported in this study were also reported to be under selection in the horse by Orlando *et al*. [3]. However, unlike our study, Orlando *et al*. [3] aimed to detect selection signatures in modern horses and included several genomes and compared ancient horse genomes to those of Przewalski’s and modern domesticated horses. Namely, the three genes shared between both studies were *NINJ1* and *SEC63* in the Tennessee Walking Horse and *COMMD1* in the Arabian horse. *NINJ1* encodes the ninjurin protein which is highly expressed in human brain endothelium [52] and becomes up-regulated after nerve injuries in Schwann cells and in dorsal root ganglion neurons [53]. *SEC63* encodes a highly-conserved membrane protein important for protein translocation in the Endoplasmic Reticulum [54]. *COMMD1* (Copper Metabolism MURR1 Domain 1) is involved in copper ion homeostasis and sodium transport [55]. It plays and essential role in the termination of NF-κB activity and control of inflammation [55,56]. *COMMD1* causes copper storage disorder in Bedlington terriers, an autosomal recessive disorder that causes rapid accumulation of copper in the liver of affected dogs [57].

### Data Availability

The raw fastq reads for all horses analyzed in the current study are available in the European Nucleotide Archive under study number PRJEB9799 (http://www.ebi.ac.uk/ena/data/view/PRJEB9799). The aligned BAM files, detected SNPs and INDELs, as well as CNVs and SVs and their functional annotations are available for download at https://www.animalgenome.org/repository/horse/6_horse-breeds_variants/. We formatted our SVs into two separate tracks, one each for inter and intra chromosomal translocations. Different colors designate different SV types in a manner similar to the system used by the human DGV [58]. Clicking on an interchromosomal translocation feature links the user to the chromosomal address of the other end of the feature. For the intrachromosmal translocations, the putative breakpoints of the feature are displayed joined together in a GFF-like style. The CNVs were formatted into a bed format with different colors for gains, losses and normal copy numbers.

## Materials and Methods

### DNA Collection and Whole Genome Sequencing

All animal procedures for DNA sampling were approved by the Cornell University Institutional Animal Care and Use Committee (protocol 2008-0121). DNA was extracted from blood using Puregene whole-blood extraction kit (Qiagen Inc., Valencia, CA, USA) or hair samples using previously published methods [59]. Paired-end sequencing was performed at the Biotechnology Resource Center, Cornell University. For the construction of sequencing libraries, genomic DNA was sheared using a Covaris acoustic sonicator (Covaris, Woburn MA) and converted to Illumina sequencing libraries by blunt end-repair of the sheared DNA fragments, adenylation, ligation with paired-end adaptors, and enriched by PCR according to the manufacturer’s protocol (Illumina, San Diego CA). The size of the sequencing library was estimated by capillary electrophoresis using a Fragment Analyzer (AATI, Ames IA) and Qubit quantification (Life Technologies, Carlsbad CA). Cluster generation and paired-end sequencing on Illumina HiSeq instruments were performed according to the manufacturer’s protocols (Illumina, San Diego) at the Biotechnology Resource Center, Cornell University. The Percheron (PER), Miniature and Arabian horse (AMH) had a library read length of 100 bp and an average insert size of 188 bp, 181 bp and 181 bp respectively. The Brazilian Mangalarga Marchador (MM), a Native Mongolian Chakouyi (CH) and a Tennessee walking horse (TWH) had a library read length of 140 bp and an average insert size of 248 bp, 168 bp and 207 bp respectively.

### Read Filtering and Alignment

A diagram summarizing the process of reads filtering, mapping, variant identification and the tools used in each step is shown in (**Figure S2)**. Raw reads were first inspected using the quality control program FastQC v10.1 (http://www.bioinformatics.babraham.ac.uk/projects/fastqc/). Then, the reads were quality filtered and adaptors removed using Trimmomatic [60]. The quality filtering utilized a sliding window of 4 bp and required a minimum mean Phred quality score of 20 within each window. Windows with an average quality less than 20 were sequentially removed from a read. Subsequently, reads with less than 60 bp of sequence remaining were removed from analysis along with their corresponding pairs. The genomes were then aligned to EquCab2 using BWA [61] in the *bwa aln* procedure and the aligned .sai files were combined into Sequence Alignment (SAM) files using *bwa sampe* procedure designated for paired end sequences. The SAM files were sorted and converted to Binary Alignment (BAM) files using PICARD toolkit v1.89 (http://sourceforge.net/projects/picard/) using *SortSam.jar*, then, the duplicate reads in the BAM file were removed using *MarkDuplicates.jar* in the same toolkit. The Genome Analysis Toolkit (GATK) v2.6-5 [62], procedures *RealignerTargetCreator* and *IndelRealigner*, were used to perform local realignment of the BAM file reads around the INDELs in order to correct misalignments due to the presence of INDELs.

### Base Quality Score Recalibration and Calling SNPs and small INDELs

SNPs and small INDELs (<50bp) were detected using the GATK *HaplotypeCaller* procedure [63]. The GATK HaplotypeCaller was designed to be very permissive so that it did not miss rare variants. In order to recalibrate base quality scores we used the *BaseRecalibrator* procedure in GATK. Since we do not currently have a gold standard set of variants for the horse (required by the procedure), we undertook an iterative approach (described in the GATK best practices (version 2.4-3). The approach simultaneously recalibrated base quality scores and eventually resulted in the final set of reported variants. First, the GATK *HaplotypeCaller* procedure [63] obtains an initial set of variants subsequently used to recalibrate base quality scores and generate recalibrated BAM files for each genome. The recalibrated files were then used to call variants in the next iteration. Subsequently, variants called in each iteration were used as a bootstrap set in place of “gold standard” variants to recalibrate the base quality scores in the following iterations. The procedure was iterated until the number of variants and the base quality score recalibrations stabilized, which in our pipeline occurred following the fifth iteration. After that, we used the GATK *VariantRecalibrator* procedure to recalibrate the variants using polymorphisms obtained from the horse genome assembly project as training set (www.ncbi.nlm.nih.gov/projects/SNP/). The VariantRecalibrator algorithm is designed to assign probabilities and quality scores used to filter out those false positives using a statistical machine learning approach. The algorithm learns the best quality score filters based on the data itself and allows the user to trade off sensitivity and specificity. It builds a Gaussian mixture model which uses variants from the input set that overlap variants in the training set. Once the model is trained, variants in the input set that have desired properties as determined by the Gaussian mixture model were filtered using the *ApplyRecalibration* procedure. After careful examination of the tranches plot (resulting from *ApplyRecalibration*) a tranches filter level of 99 was used as it resulted in the highest number of true positive SNPs while minimizing false positives (**Figure S3**).

### Identifying Structural Variations and Copy Number Variations

Structural variations (SVs) and large INDELs were identified using SVDetect [64]. The program uses anomalously mapped read pairs to localize rearrangements within the genome and classify them into their various types. After filtering out correctly mapped pairs, we used a sliding window of size 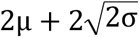 to partition the genome, where *μ* is the estimated insert size and *σ* is the standard deviation. The length of steps in which the sliding window moved across the genome were set to half of the window size. Control-Freec [65] was used to detect copy number variations (CNVs). The program uses GC-content and mapability profiles to normalize read count and therefore gives a better estimate of copy number profiles in high GC or low coverage regions [65]. A breakpoint threshold of 0.6 and a coefficient of variation of 0.05 were used in the analysis.

### Variant Annotation

We used SNPEff v4.0 [66] to annotate the SNPs and short INDELs using the latest available ENSEMBL gene annotation database (EquCab2.76). The output of SNPEff is a full list of effects per variant. SNPs and Indels located within 5,000 bases (5 kb) upstream or downstream genes as well as those within exons, introns, splice sites, and 5’ and 3’ untranslated regions (UTRs) were also annotated. Since SNPEff output can be integrated into GATK VCF file, we have produced an annotated version of the GATK VCF file which can be loaded and viewed easily in genome browsers. The CNVs and SV breakpoints overlapping ENSEMBL genes were detected using Bedtools (v2.23.0). Ensembl gene IDs were then converted to gene names using Biomart.

We used Nucleotide Diversity (π) to identify regions with low diversity. For each of the genomes, the nucleotide diversity (π) was calculated for the SNPs in 1 MB non-overlapping windows using VCFtools v1.10 [50]. Regions in the lower 1% tail of the π distribution were considered low diversity regions following a similar approach by Branca *et al.* [67]. Genes in these regions were annotated for biological process, using PANTHER v14.0 [68]. Circos plots [69] summarizing the distribution of the genomic variations were then created for each of the genomes and a summary circos plot was created to highlight variants in common between the six genomes. To enhance visualization, we removed the small intrachromosomal elements (endpoints size <10 bp) and interchromosomal elements (endpoints size <500 bp) due to their abundance in the output which makes it difficult to visualize in the circos plot.

### RT q-PCR analysis of the Latherin CNV

We used Quantitative PCR to quantify the copy number variation in exons of 4 genes from 8 horses, including the horse used to produce the EquCab 2.0 reference genome assembly. Primers designed in Primer3 v0.4.0 targeted exons overlapping the NGS identified copy number variation [70] (**Table 3**). Genomic DNA (25 ng) was amplified in 10 uL reactions using the Quanta Biosciences PerfeCta SYBR Green (FastMix) as per the manufacturers recommended conditions (Gaithersburg, MD, USA). *ASIP* exon 2 was amplified as reference single-copy gene. Thermocycling and detection were performed using PCR on the Illumina Eco Real-Time PCR System using parameters recommended for the Quanta Mix (58°C annealing). Copy numbers were calculated relative to the reference genome horse. We substituted the Percheron and American Miniature horses by horses from the same breed, as DNA samples from the original two sequenced horses were unavailable.

**Table 3:**
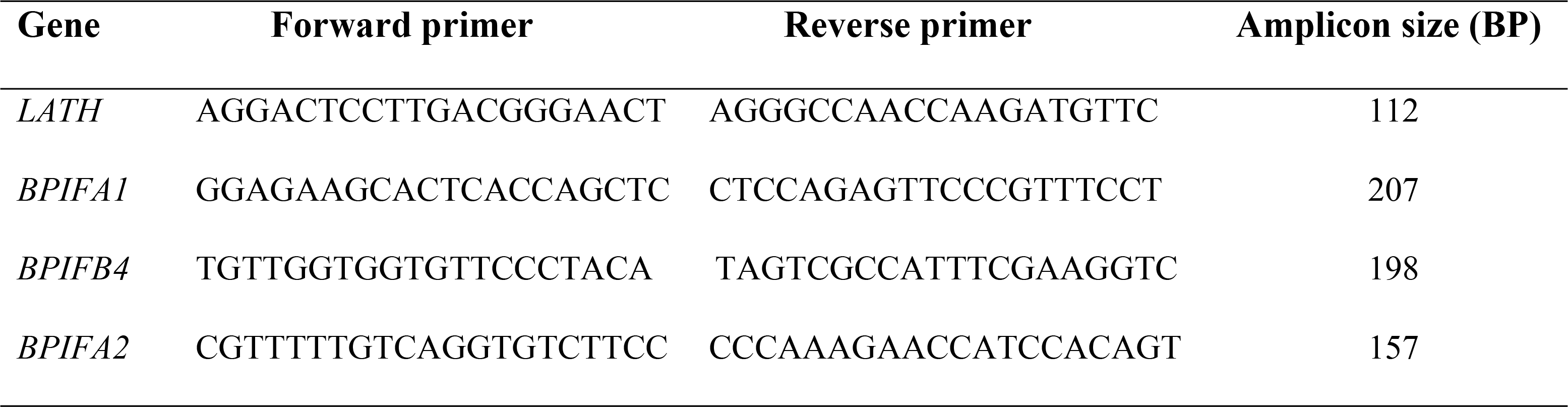
Real-time quantitative PCR primers.

## Acknowledgements

The authors would like to thank all owners who volunteered samples from their horses for the study. The authors would also like to thank Chuzhao Lei, Yun Ma and Wansheng Liu for providing the DNA of the Native Mongolian Chakouyi horse. The authors also thank Dr. Doug Antzack and Dr. Donald Miller for providing part of the computational resources based on which part of this work was completed.

## Supporting Information

**Table S1:** Annotation of SNPs and INDELs by putative functional consequence.

**Table S2:** Annotations of Structural variants and SVs and Copy number variants (CNVs) detected in six horses. ENSEMBL gene names and their corresponding biomart IDs (when available) overlapping the variants locations are given.

**Table S3**: Chromosomal regions and genes in high nucleotide diversity (π) regions (top 1% of the empirical distribution).

**Table S4**: Statistical overrepresentation test (Bonferroni-corrected for P < 0.05) for genes in high π regions in the six horses in chromosomes 12 and 20.

**Table S5**: Chromosomal regions and genes in low high nucleotide diversity (π) regions (lower 1% of the empirical distribution).

**Table S6**: Statistical overrepresentation test (Bonferroni-corrected for P < 0.05) for genes in low π regions in various horses.

**Table S7:** PANTHER GO-Slim biological processes for xenoRefGene genes in regions with low pi values in these 6 horse genomes. The table shows, for each horse, the percentage of genes hit against total number of processes hits. The average for each biological category shows abundance in metabolic processes and biological regulation compared to other. Genes pertaining to growth are present in the Percheron and American miniature horses only and cell proliferation is only present in the Arabian horse.

**Figure S1:** The allele frequencies of SNPs from whole-genome sequence data of the six horses showing a lower frequency as observations of the alternate allele increased.

**Figure S2:** An overview of the pipeline used in the reads processing and variant detection. Description of the step and the name of the program/software given in parenthesis.

**Figure S3:** Tranches plot generated by GATK VariantRecallibration procedure. The plot shows the trade off in (gain in cumulative false positives (FP)) resulting from choosing a certain level of cumulative true positive (TPs) variants. Tranche specific true positives and false positives are shown in blue and orange respectively.

